# Human CTC1 primarily functions in telomere maintenance/protection and promotes CHK1 phosphorylation in response to global replication stress

**DOI:** 10.1101/2020.01.15.906891

**Authors:** Stephanie M. Ackerson, Caroline I. Gable, Jason A. Stewart

## Abstract

CST (CTC1-STN1-TEN1) is a heterotrimeric, RPA-like protein that binds to single stranded DNA (ssDNA) and functions in the replication of telomeric and non-telomeric DNA. Previous studies have shown that deletion of CTC1 results in decreased cell proliferation and telomeric DNA damage signaling. However, a detailed analysis of the consequences of conditional CTC1 knockout (KO) have not been fully elucidated. Here, we investigated the effects of CTC1 KO on cell cycle progression, genome-wide replication and activation of the DNA damage response. We find that CTC1 KO results in p53-mediated G2 arrest and increased apoptosis, but not genome-wide replication defects or DNA damage. Instead, the G2 arrest is dependent on the accumulation of telomeric RPA following CTC1 KO, suggesting that the primary function of CST is in telomere end protection and maintenance not genome-wide replication. However, despite increased RPA-ssDNA, global CHK1 phosphorylation was not detected in CTC1 KO cells. Further analysis revealed that CTC1 KO significantly inhibits CHK1 phosphorylation following hydroxyurea-induced replication stress, due to decreased levels of the ATR activator TopBP1. Overall, our results identify that telomere not genome-wide DNA damaging signaling leads to decrease proliferation following CTC1 deletion and that CST promotes ATR-CHK1 signaling through the regulation of TopBP1.

## INTRODUCTION

DNA damage can arise from both endogenous and exogenous factors, such as replication stress, ionizing radiation and oxidation. To prevent genome instability and disease, cells have evolved elaborate signaling pathways to sense the damage, arrest the cell cycle and repair the DNA. This process, known as the DNA damage response (DDR), is primarily mediated by three members of the Phosphatidylinositol-3-kinase-related kinase (PIKK) family known as ataxia telangiectasia mutated (ATM), ataxia telangiectasia and Rad3-related (ATR), and DNA-dependent protein kinase (DNA-PK) (1). Activation of these kinases in response to DNA damage leads to the phosphorylation of downstream effectors and stabilization of the tumor suppressor, p53. p53 in turn induces transcription of downstream targets, such as p21 and p16, that facilitate cell cycle arrest and eventual apoptosis or senescence, if the damage is not resolved.

ATM and DNA-PK are primarily activated in response to double-strand breaks (DSBs), whereas ATR coordinates the repair of DNA damage arising from single-stranded (ss)DNA gaps or breaks. ATR plays a primary role in managing replication stress during S-phase and is essential for the survival of dividing cells (2-4). During S-phase, ssDNA can arise from uncoupling of the helicase from the replisome and nucleolytic processing of various replication and repair intermediates (5,6). Once formed, large regions of ssDNA are quickly recognized and bound by the ssDNA binding protein, replication protein A (RPA) (7). ATR interacting protein (ATRIP) then associates with the RPA-bound ssDNA (RPA-ssDNA), which localizes ATR (8). Localization of ATR, however, is not sufficient to fully activate ATR kinase activity.

In vertebrates two major ATR activating proteins have been identified, topoisomerase 2-binding protein 1 (TopBP1) and Ewing’s tumor-associated antigen 1 (ETAA1) (9-12). Both proteins contain ATR activating domains (AAD) that modulate ATR kinase activity (13). ETAA1 was only recently discovered and less is known about the mechanism by which it activates ATR. However, recent work suggests that ETAA1 plays only a minor role in ATR activation during the DDR (14,15). Instead, ETAA1 plays a primary function in proper chromosome alignment and checkpoint activation in metaphase as well as preventing untimely entry into G2. On the other hand, TopBP1 is essential for ATR activation in response to ssDNA damage (12). For ATR activation through TopBP1, the RAD9-RAD1-HUS1 (9-1-1) complex is loaded at ssDNA-dsDNA junctions independent of ATRIP-ATR recruitment to RPA-ssDNA (16,17). TopBP1 is then recruited to the 9-1-1 complex and interacts with ATR-ATRIP, stimulating ATR kinase activity. Once activated, ATR phosphorylates numerous downstream targets, including checkpoint kinase 1 (CHK1) and p53 (18-20). CHK1 then promotes the degradation of CDC25A, leading to inactivation of cyclin-dependent kinases and inhibition of replication origin firing (21). Failure to activate the ATR-CHK1 pathway, particularly following treatment with replication inhibitors or in cancer cells with high levels of genome instability, leads to impaired growth and cell death. In fact, several ATR/CHK1 inhibitors are currently in clinical trials or under development as cancer therapeutics (22).

In the current study, we examined the effects of conditional gene knockout of the largest subunit of CST (CTC1-STN1-TEN1), CTC1, on cell growth and checkpoint activation. The mammalian CST complex is a conserved ssDNA binding protein with structural similarities to RPA and interacts with DNA polymerase α-primase (pol α) (23). Like RPA, CST has multiple oligosaccharide-oligonucleotide binding folds (OB-folds) that allow dynamic binding to ssDNA. CST plays a critical role in telomere replication and has recently been shown to function in genome-wide replication and DSB repair (24-30). At telomeres, CST promotes telomerase dissociation following telomere extension and is then required to convert the G-rich ssDNA overhang (G-overhang) to duplex DNA through a process known as C-strand fill-in (31,32).

Previous work demonstrated that conditional deletion of human CTC1 leads to a significant growth delay and unregulated lengthening of the G-overhang by telomerase as well as shortening of telomeres from the inability to perform C-strand fill-in (33). G-overhangs are typically protected by POT1, a member of the shelterin complex. However, these extended G-overhangs exhaust the cellular pools of POT1, leading to telomeric RPA-ssDNA and telomeric γH2AX, a marker of DNA damage. While this accumulation of RPA-ssDNA is predicted to elicit activation of the ATR-CHK1 pathway, this was not directly tested in previous studies (33,34). Furthermore, it was unclear whether the growth inhibition following CTC1 deletion was primarily due telomere or non-telomere defects. Here, we performed a detailed cell cycle analysis following conditional CTC1 deletion to determine how it affected cell cycle progression and the DDR. Our findings indicate that the growth inhibition caused by CTC1 removal is dependent on the accumulation of telomeric RPA-ssDNA and not defects in genome-wide DNA replication or DNA damage. Moreover, we were surprised to find that, while CTC1 deletion activates ATR, we were unable to detect CHK1 phosphorylation, even though RPA-ssDNA and γH2AX were present. In addition, we found that ATR-mediated CHK1 activation is significantly inhibited in response to HU-induced fork stalling, due to decreased levels of TopBP1. Overall, this work provides important insights into the telomere versus non-telomere functions of CTC1 and implicate CST as a novel regulator of the ATR-CHK1 pathway.

## MATERIALS AND METHODS

### Cell Culture

HCT116 cells were maintained in McCoy’s 5A media supplemented with 10% fetal bovine serum and 1% penicillin/streptomycin at 37°C with 5% CO2. Cells were checked regularly for mycoplasma contamination. HCT116 CTC1^F/F^, CTC1^F/F^+Flag-CTC1 and CTC1^F/F^+Flag-POT1 cells were generously provided by Dr. Carolyn Price (33).To induce Cre-ER mediated recombination of the CTC1 gene, a final concentration of 10 nM tamoxifen (TAM) was added to CTC1^F/F^ cells. The initial addition of TAM is indicated as day 0. At each passage, 10 nM TAM was again added to ensure CTC1 gene disruption. For siRNA p53 knockdown, 5 nM ON-TARGETplus siRNA SMART pools (Dharmacon) to p53 (L-003329-00) or non-targeting control (D-001810-10-05) were transfected into cells with Lipofectamine RNAiMAX (Thermo Fisher Scientific). For TopBP1 transfection, cells were plated 24 h before transfection in 100 mm dishes at 1×10^6^ cells. 12.5 μg of pcDNA3-TopBP1 was mixed with 25 μl of Polyethylenimine (10mg/ml) (Polysciences) in a total volume of 550 μl for each transfection. The pcDNA3-TopBP1 plasmid was generously provided by Dr. Weei-Chen Lin (35). After 48 h, the cells were collected and Whole Cell Protein Extraction performed (see below).

### Whole Cell Protein Extraction

Cell pellets were lysed, sonicated and nuclease-treated as previously described (25). The supernatant was collected and protein concentration measured by BCA assay. The samples were then mixed with SDS-PAGE loading buffer and analyzed by western blotting as described below.

### Subcellular Fractionation for Protein Extraction

Cell pellets were lysed in 200 µL Buffer A (10 mM HEPES pH 7.9, 10 mM KCl, 1.5 mM MgCl_2_, 0.34 M Sucrose, 1 mM DTT, 0.1% Triton X-100, 1x phosphatase inhibitors [4 mM β-glycerophosphate, 4 mM sodium vanadate, and 20 mM sodium fluoride] and 1x protease inhibitors [1 µg/mL pepstatin A, 5 µg/mL leupeptin, 1 µg/mL E64, 2 µg/mL aprotinin, and 5 µg/mL antipain]) and incubated on ice for 10 min. Cell lysates were centrifuged at 4°C 1300 *× g* for 5 min. The supernatant was transfered to a new tube and excess cell debris was removed by centrifugation at 20,000 *× g* for 15 min at 4°C. The supernatant (Cytosolic Fraction) was transfered to a new tube. The cell pellet containing nuclei was resuspended in 100 µl Buffer B (3 mM EDTA pH 8.0, 0.2 mM EGTA, 1 mM DTT, 1x phosphatase inhibitors, and 1x protease inhibitors) and were incubated on ice for 30 min with mixing at 15 and 30 min. The samples were then centrifuged at 2,000 *× g* for 5 min at 4°C and the supernatant containing the soluble nuclear fraction was transferred to a new tube. The cell pellet containing the chromatin bound fraction was resuspended in 100 µL Buffer A and then sonicated at 40% amplitude for 3 cycles of 10 seconds on and 5 seconds rest. Then, samples were treated with Benzonase (0.0625 U/ µL; EMD Millipore) for 1 h on ice. The samples were then centrifuged at 15,800 *× g* for 10 min at 4°C and the supernatant saved as the chromatin fraction. Protein concentrations were determined with the BCA assay and samples analyzed by western blot as described below.

### Western Blot Analysis

20-40 µg of protein were run by SDS-PAGE and transferred to a nitrocellulose membrane. All membranes were checked with Ponceau S staining for transfer efficiency and total protein loading. Membranes to analyze CTC1 levels were blocked with 3% BSA in 1x PBS plus 0.1% Tween 20 (PBST) for at least 2 h, and all subsequent antibodies were diluted in 3% BSA-PBST. For analysis of CHK1 S317, membranes were blocked in 5% non-fat milk in 1x TBS plus 0.1% Tween 20 (TBST) for at least 2 h, and all subsequent antibodies were diluted in 5% non-fat milk-TBST. For all other western blot analysis membranes were blocked in 5% non-fat milk-PBST for at least 2 h. Primary antibodies were diluted in 5% non-fat milk-PBST and incubated at 4°C overnight. Primary antibodies were removed and the membranes washed 3 × for 10 min each in PBST (TBST for CHK1 S317 analysis). Secondary antibodies were diluted in the solution indicated above for at least 2 h at room temperature. After incubation the blots were then developed with Western Lightning Plus ECL (Perkin Elmer) or ECL Prime (GE Healthcare).

### Flow Cytometry

The cells were collected and washed with 1x PBS. After the supernatant was removed, 5 mL of ice-cold 100% methanol was added drop wise with gentle vortexing. Tubes were then placed at −20°C for 10 min followed by centrifugation at 2000 *× g* for 5 minutes. The supernatant was removed and the cell pellets were washed with 5 mL 1x PBS and centrifuged again at 2000 *× g* for 5 minutes. The supernatant was then removed and the cell pellets were stored at 4°C at least overnight.

To detect S-phase cells, EdU (50 μM) was added 30 min prior to collection. EdU was detected by Click-iT chemistry, according to the manufacturer’s instructions (Thermo Fisher). Cells were resuspended in 250 µL of Click-iT reaction cocktail and incubated for 30 minutes at RT protected from light. 5 mL of 1% BSA-PBST was then added and samples were spun down at 1000 x *g* for 5 minutes and the supernatant was removed. The cells were then resuspended in 650 µL of fresh DAPI Staining Solution (0.1% Triton X-100, 0.1 mg/mL RNase, 1 µg/mL DAPI diluted in 1 × PBS). The samples were spun at 50 *× g* for 30 s to remove cell clumps and debris through filter-capped tubes (Corning) and run on a BD LSR II Flow Cytometer in the Microscopy and Flow Cytometry Facility at the University of South Carolina, College of Pharmacy.

### Immunofluorescence (IF) and IF-fluorescence in situ hybridization (IF-FISH)

Cells were plated onto coverslips and allowed to grow for 24 hours to 50-70% confluency. 50 µM EdU was added for 30 minutes prior to collection. For detection of phosphorylated H2AX S140 (γH2AX), Histone H3 Ser10, and RPA32 Ser33, cells were fixed with 4% formaldehyde in 1 × PBS for 10 min at RT. After formaldehyde incubation, cells were rinsed twice with 1x PBS and then permeabilized with 0.5% Triton X-100 diluted in 1 × PBS for 10 min at RT. Slides were washed with 1 × PBS then stored in 1 × PBS at 4°C overnight. For detection of chromatin bound RPA32, cells were pre-extracted with 0.1% Triton X-100 in 1 × CSK buffer for 5 minutes at room temperature. Slides were then washed once with 1 × PBS and fixed by adding 1 mL of 100% ice cold methanol. Slides were incubated at −20°C for 10 minutes then washed with 1 × PBS then stored in 1 × PBS at 4°C overnight. IF was then performed as previous described for γH2AX (1:5000), phosphorylated Histone H3 S10 (1:500), phosphorylated RPA32 S33 (1:1000) or RPA32 (1:500) and nuclear intensity measured in ImageJ as previously described (25).

For IF-FISH, IF was performed for γH2AX or chromatin-bound RPA32, as described above. Telomere FISH was then performed, as previously described (36). Briefly, after the last wash, following secondary antibody incubation, the coverslips were fixed with 2% formaldehyde in 1x PBS for 10 min at RT. After two washes with 1x PBS, the coverslips were dehydrated and incubated with a telomeric G-strand PNA probe (AlexaFluor 488-TTAGGG_3_; PNA Bio) in hybridization buffer (10 mM Tris pH 7.4, 70% formamide, 1% blocking reagent [Roche]) for 10 min at 80°C. Coverslips were then incubated at RT for 2-3 h followed by two washes in wash buffer (10 mM Tris pH 7.4, 70% formamide) for 15 min each. The coverslips were subsequently washed three times with 1x PBS, dehydrated and mounted on slides with mounting media containing 0.2 μg/ml DAPI. IF and IF-FISH images were then taken under 40x or 60x on an EVOS epifluorescence microscope (Thermo). Signal intensities and foci were then scored using ImageJ.

### Antibodies and Chemical Inhibitors

Primary: CTC1 (33), STN1 (Abcam, 119263), Actinin (Santa Cruz, sc17829), α-Tubulin (Sigma, T-9026), CHK1 (Bethyl, A300-298A), CHK1 S317 (Bethyl, A304-673A), p53 (Cell Signaling, 9282), p21 (Santa Cruz, sc-6246), H3 (Cell Signaling, 9715), pH3 S10 (Cell Signaling, 9706), Rad9 (Santa Cruz, sc-74464), RPA32 (Abcam, ab16850), pRPA32 S33 (Bethyl, A300-246A), TopBP1 (Bethyl, A300-111A), ETAA1 (kindly provided by Dr. David Cortez), γH2AX (Bethyl, A300-081A) and POT1 (Abcam, ab124784). Secondary: Thermo: anti-rabbit-HRP (32460); anti-mouse-HRP (32430); Molecular Probes: goat-anti-rabbit AlexaFluor 647 (A21244), goat-anti-mouse AlexaFluor 647 (A21235), goat-anti-mouse AlexaFluor 594 (A11032), goat-anti-rabbit AlexaFluor 594 (A11037). ATR inhibitor: VE-821 5 µM for 24 h (Selleckchem, S8007).

## RESULTS

### CTC1 deletion leads to decreased proliferation and increased G2 cells

To more closely examine the effects of CTC1 gene disruption, we performed time course analysis of cell cycle progression in HCT116 conditional CTC1 knockout (CTC1^-/-^) cells. This cell line was previously generated by the addition of loxP sites surrounding exon 5 of both endogenous *CTC1* alleles (CTC1^F/F^) (33). Cells were then stably selected for expression of a Cre recombinase linked to the estrogen receptor (Cre-ER). Addition of tamoxifen (TAM) results in localization of Cre-ER to the nucleus and removal of exon 5 by Cre-induced recombination. Gene disruption was confirmed by Western blot and PCR analysis following the addition of TAM, as previously described (Figure 1A and S1A) (33). We observed that endogenous CTC1 protein levels significantly decreased after three days and were almost completely absent four days after TAM addition (Figure 1A). To control for possible off-target effects, a stable cell line was developed, expressing Flag-CTC1 in the CTC1^F/F^ line (CTC1^F/F^+Flag-CTC1) (33). Following the addition of TAM to this cell line (CTC1^-/-^+Flag-CTC1), the endogenous CTC1 is disrupted, while the exogenous Flag-CTC1 expression remains unchanged (Figure 1A and S1A).

**Figure 1:**
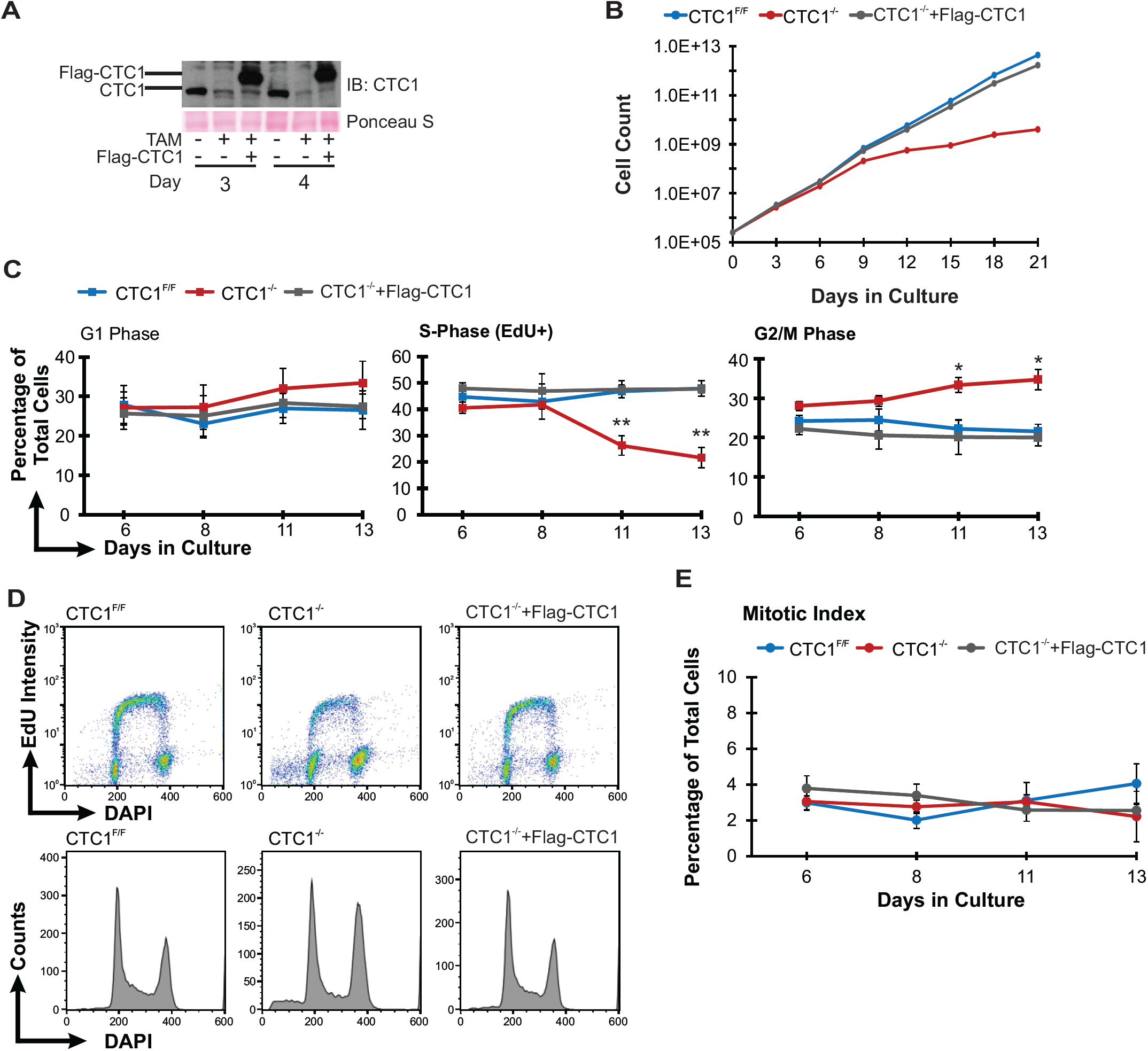
CTC1 deletion results in decreased cell proliferation. **(A)** Western blot of CTC1 knockout in HCT116 cells. Tamoxifen (TAM) was added to CTC1^F/F^ and CTC1^F/F^+Flag-CTC1 cell lines at day 0 to disrupt the CTC1 gene (CTC1^-/-^). Representative gel showing days 3 and 4. Ponceau S staining is used as a loading control. **(B)** Representative growth curve of three independent biological trials. **(C-D)** Flow cytometry analysis of HCT116 CTC1^F/F^, CTC1^-/-^, and CTC1^-/-^ +Flag-CTC1 cells. **(C)** Percentage of cells in each cell cycle phase, as indicated. (n=3, independent biological experiments). **(D)** Representative histograms from flow cytometry analysis. Top Panel: DNA content (DAPI) versus replicating cells (EdU+). Bottom Panel: DAPI versus cell count. **(E)** Mitotic index is based on the levels of phosphorylated Histone H3 as measured by immunofluorescence (n=3, independent biological experiments). Error bars indicate ±SEM. P-values were calculated by an unpaired, two tailed *t* test (\**P* ≤0.05, \*\**P* ≤0.01).

To better understand the effects of CTC1 removal on cell proliferation, growth curve and cell cycle analysis were performed (Figure 1B-D). In agreement with previous findings, we observed decreased cell proliferation starting around 6 days after conditional CTC1 deletion, which became more severe at 10-12 days in culture (Figure 1B). We expected that the growth delay was caused by an intra-S-phase checkpoint triggered by replication issues previously observed after depletion of CST subunits (25,29,30,37). However, while cell cycle analysis revealed a substantial decrease in the number of S-phase cells following CTC1 deletion, the cells within S-phase retained similar levels of DNA synthesis (EdU intensity per cell) compared to controls (Figure 1C-D), suggesting that replication is not inhibited in the CTC1^-/-^ cells. In order to confirm this result, we performed DNA fiber analysis on day 11 after the TAM addition (Figure S1B-D). In agreement with our flow cytometry data, we failed to detect any significant changes in DNA synthesis or replication events following CTC1 deletion. Unexpectedly, these findings indicate that global replication rates are not significantly altered following CTC1 removal. Instead, we observed a significant accumulation of G2/M cells starting at day 11 after TAM addition (Figure 1C). These changes in cell cycle profile mirror the decrease in cell proliferation observed in the CTC1^-/-^ cells (Figure 1B), suggesting a delay or arrest in cell cycle progression.

In order to distinguish between G2 versus M-phase, we used immunofluorescence (IF) to assess the number of phosphorylated Histone H3 S10 positive cells, as a readout of the mitotic index (Figure 1E) (38). The percentage of mitotic cells was not increased in the CTC1^-/-^ cells compared to controls, indicating that CTC1 deletion causes an accumulation of G2 rather than M-phase cells. In addition, we observed a significant increase in the sub-G1 population (Figure 1D), which could arise from increased apoptosis. Indeed, we found that CTC1 deletion led to increased caspase 3/7 activity (Figure S1E). Together, these results indicate that CTC1 deletion leads to an accumulation of G2 arrested cells and increased apoptosis. A previous study also showed that conditional CTC1 deletion leads to increased senescence, suggesting that both apoptosis and senescence contribute to overall growth inhibition in CTC1^-/-^ cells (33).

### CTC1 KO induces a p53-dependent DDR but not CHK1 phosphorylation

Since changes in global replication were not observed, we expected that growth inhibition and increased cell death are due the gradual accumulation of DNA damage at telomeres and other G-rich sites across the genome, which have shown fragility with CTC1 or STN1 depletion (34,39). To determine whether the growth inhibition in the CTC1^-/-^ cells was mediated by p53, we next measured the levels of p53 and p21. Cells were collected at days 8, 11 and 13 after TAM addition and whole cell extracts analyzed by Western blot (Figure 2A). Both p53 and p21 were significantly increased starting at eight days after TAM addition, which corresponds to the partial G2 arrest and growth inhibition in the CTC1^-/-^ cells (Figure 2A). In order to confirm that the partial G2 arrest was dependent on p53, CTC1^-/-^ cells were treated with siRNA to p53 eight days after TAM addition, collected on day 11 and cell cycle analysis performed. siRNA depletion of p53 largely rescued the G2 arrest (Figure S2A-B). The incomplete rescue is likely due to incomplete knockdown of p53, which resulted in low levels of p21 expression (Figure S2A). Our findings indicate that CTC1 deletion leads to a p53-mediated G2 arrest.

**Figure 2:**
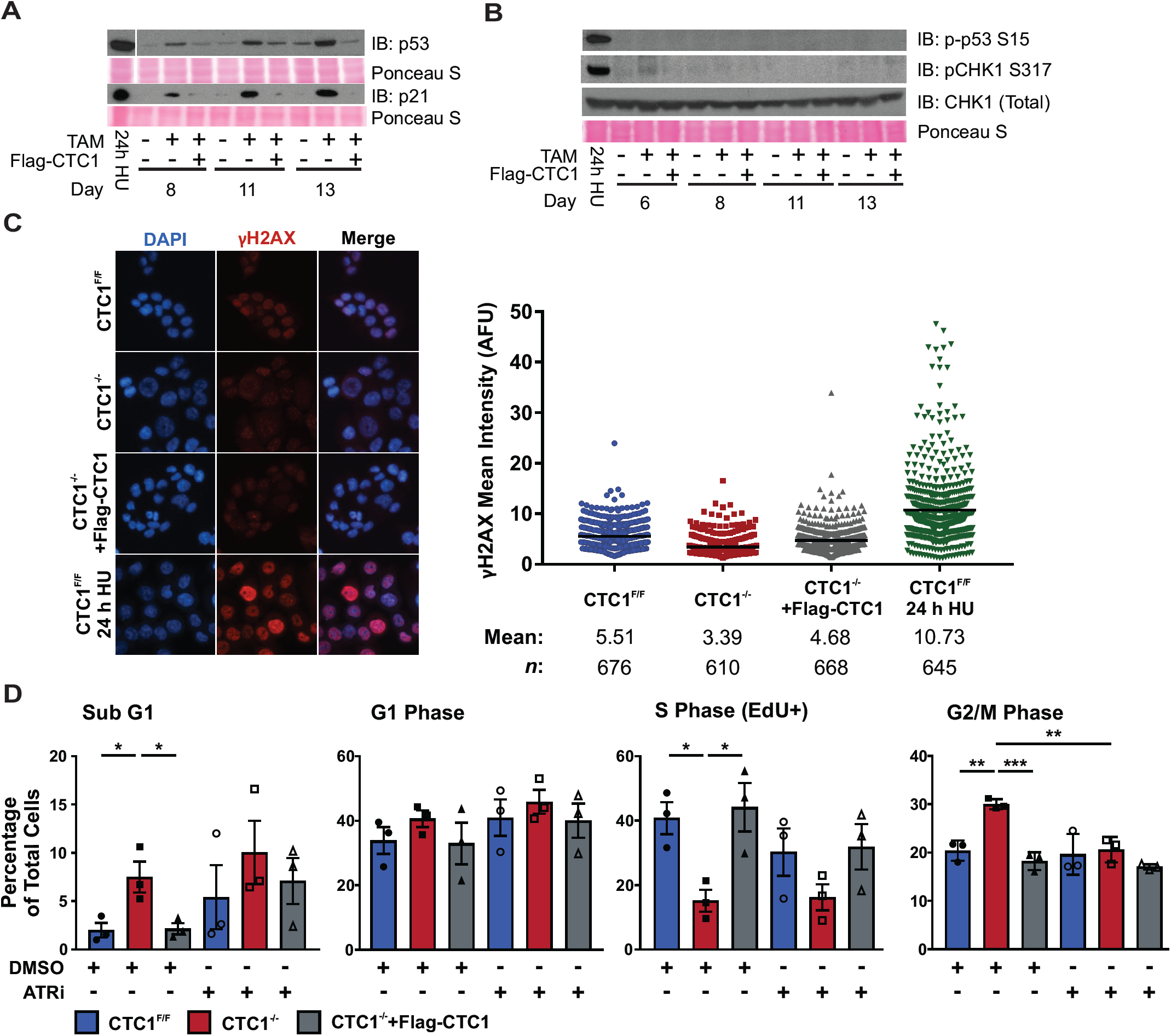
CTC1 deletion induces a p53 dependent cell cycle arrest through ATR signaling and not CHK1 activation. **(A-B)** Western blot of whole cell extracts from HCT116 cells. Ponceau S staining is used as a loading control. Cells treated with 24 h, 2 mM hydroxyurea (24h HU) serve as a positive control. **(A)** Levels of p53 and p21. **(B)** Levels of phosphorylated p53 S15 or CHK1 S317. Total CHK1 levels serve as a loading control for pCHK1 S317 and total p53 levels are shown in (A). **(C)** Left: Representative images of γH2AX levels in HCT116 cells, as indicated. DAPI: blue, γH2AX: red. Right: Dot plots of mean γH2AX intensity per nuclei in arbitrary fluorescent units (AFU) for each nuclei. Black line and numbers below the graph indicate the mean AFU. 24 h HU sample serves as a positive control for γH2AX. Data is representative of three independent biological experiments at 13 days after addition of TAM. *n* indicates the number of total nuclei scored. **(D)** Cell cycle profile analysis at 13 days after addition of TAM in HCT116 cells treated with DMSO or 5 µM ATR inhibitor (ATRi; VE-821) for 24 h. (n=3, independent biological replicates.) Error bars indicate the ±SEM. P-values were calculated by an unpaired, two tailed *t* test (\**P* ≤0.05, \*\**P* ≤0.01, \*\*\**P* ≤0.001).

Since p53 levels were increased, we next assessed whether G2 arrest was ATR-dependent. We hypothesized that the increase in p53 was caused by activation of the ATR-CHK1 pathway following conditional CTC1 deletion because previous studies found that, in the absence of CTC1, telomerase overextends G-overhangs, leading to RPA binding and telomeric γH2AX (33). Following the accumulation of RPA-ssDNA, ATR should be recruited followed by the phosphorylation of several downstream ATR targets, including H2AX, CHK1 and p53. As a readout of ATR activation, we measured the phosphorylation of CHK1 S317 (pCHK1 S317) and p53 S15 in whole cell lysates as well as global γH2AX levels by IF. Surprisingly, we found that CTC1 KO did not result in detectable CHK1 S317 or p53 S15 phosphorylation (Figure 2B) or increased global γH2AX (Figure 2C), suggesting that the ATR-CHK1 pathway may be defective in the CTC1^-/-^ cells.

To directly assess ATR activation, we next performed flow cytometry on cells treated with the ATR inhibitor (ATRi) VE-821 for 24 h (Figure 2D and S3). Unexpectedly, we found that ATR inhibition prevents the accumulation of G2 cells following CTC1 deletion, suggesting that the arrest is ATR-dependent (Figure 2D). ATR inhibition also resulted in a further increase in apoptosis, as measured by caspase 3/7 activity in CTC1^-/-^ cells (Figure S1E). This is consistent with previous studies demonstrating that ATR inhibition increases cancer cell death, due to the loss of cell cycle checkpoints (22). These results indicate that G2 arrest is dependent on ATR kinase activity. However, this level of ATR signaling is insufficient to induce detectable levels of pCHK1 S317 (Figure 2B). Although previous work suggest that this inhibitor is highly specific to ATR versus related kinases, such as ATM and DNA-PK, it is also possible that the rescue could be due to off-target ATRi effects (40,41).

### Deletion of CTC1 leads to increased telomeric but not genome-wide RPA-ssDNA

We were surprised to find that removal of CTC1 did not result in ATR-mediated CHK1 activation, particularly given the role of CST in telomere and genome-wide replication. As mentioned previously, CTC1 KO leads to telomeric RPA and γH2AX staining when assessed by metaphase spread analysis (33). These findings indicate that RPA-bound ssDNA is generated in CTC1^-/-^ cells. However, it was possible that the amount of ssDNA was not sufficient to induce global CHK1 phosphorylation. To determine global RPA-ssDNA levels following CTC1 deletion, we used IF to measure chromatin-bound RPA levels in interphase cells (Figure 3). Cells were pre-extracted and fixed to determine the amount of nuclear RPA foci as a readout of RPA-ssDNA (Figure 3A). We found that indeed CTC1^-/-^ cells have a significant increase in cells containing RPA foci (Figure 3B). Interestingly, a subset of CTC1^-/-^ cells had enlarged nuclei, which typically associated with increased RPA-foci (i.e. larger nuclei, more RPA foci) (Figure S4). Furthermore, flow cytometry analysis was performed to determine when chromatin-bound RPA levels increased. Cells were labeled with EdU for 30 min prior to collection, pre-extracted and both chromatin-bound RPA and EdU were detected (Figure S5A and B). This analysis revealed that increased levels of chromatin-bound RPA primarily occurred in G2/M, suggesting that the accumulation of RPA-ssDNA is the primary cause of G2 arrest (Figure S5B-C).

**Figure 3:**
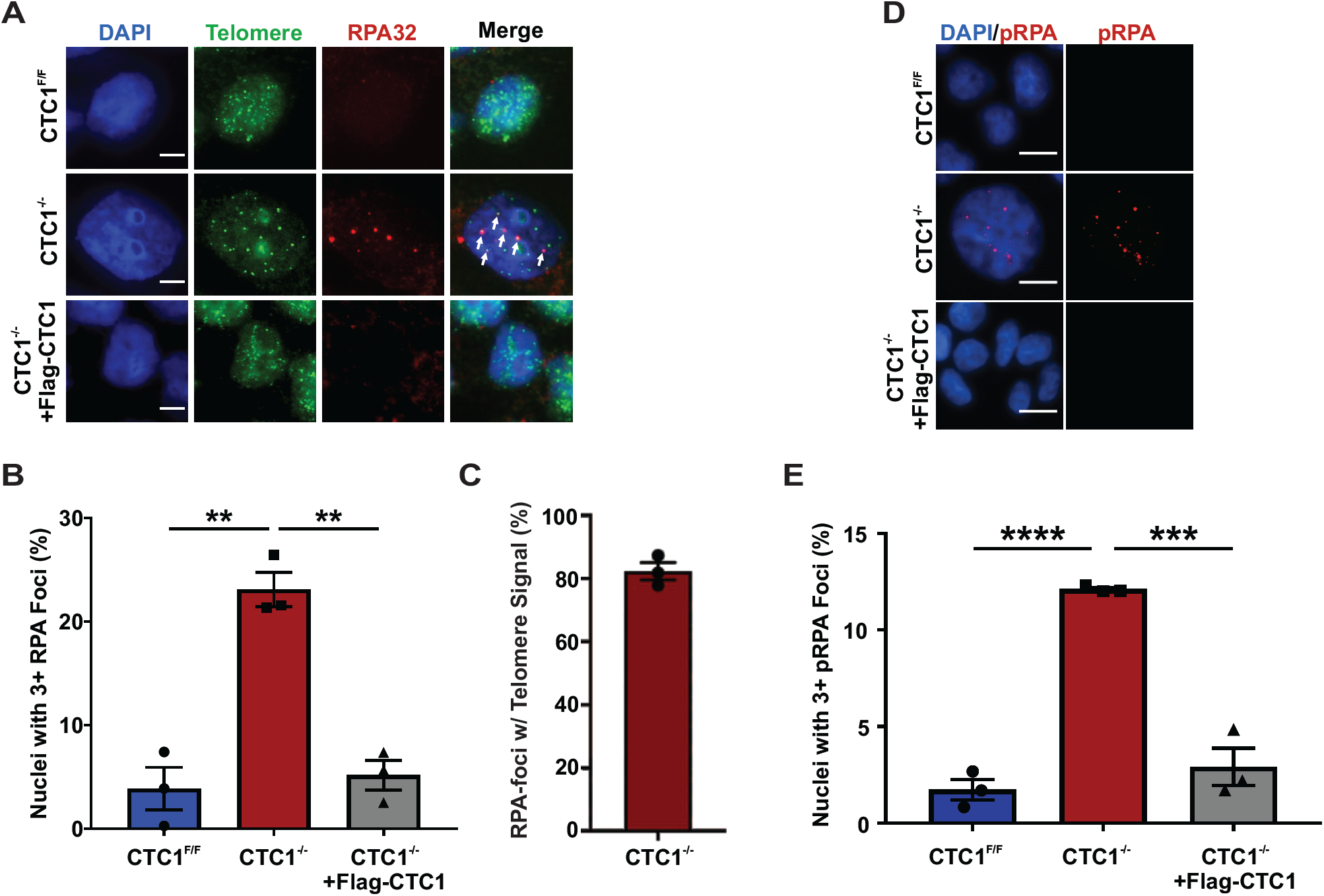
Telomeric RPA levels are increased in response to CTC1 KO. **(A)** Representative images of RPA foci and telomere FISH signal in HCT116 cells, as indicated. Cells were pre-extracted prior to fixation to identify chromatin-bound RPA. DAPI: blue, RPA32: red, Telomere (TTAGGG_3_): green. Scale bar represents 5 µm. **(B)** Percentage of nuclei with three or more RPA foci. (n=3, independent biological replicates.) **(C)** Percentage of RPA-foci colocalizing with telomere signal. (n=3, independent biological replicates.) (**D**) Representative images of phosphorylated RPA S33 (pRPA) foci in HCT116 cells, as indicated. Scale bar represents 25 µm. **(E)** Shows the percentage of cells with greater than three pRPA S33 foci. (n=3, independent biological replicates). Error bars indicate the ±SEM. P-values were calculated by an unpaired, two tailed *t* test (\*\**P* ≤0.01, \*\*\**P* ≤0.001, \*\*\*\**P* ≤0.0001).

Since previous work had shown that RPA is increased at telomeres on metaphase chromosomes following CTC1 deletion, we predicted that at least a subset of these foci were localized to telomeres (33). Yet, we predicted that they were also found at other sites across the genome. In order to distinguish between telomere and non-telomere RPA foci, we performed telomere FISH combined with IF to measure the percentage of RPA foci colocalizing with telomeric DNA (36). As expected, we observed colocalization of RPA foci with telomeres (Figure 3A and C). Unexpectedly, these RPA foci were almost exclusively at telomeres (∼80%). We suspect that the small percentage of non-telomeric RPA foci are due to the limits of telomere FISH detection, as the telomere signal was weak in many of the RPA foci. This is likely due to concurrent telomere shortening arising from defective C-strand fill-in in CTC1^-/-^ cells (33). However, it is possible that other regions of the genome may also have RPA-ssDNA in the absence of CTC1 or that RPA binding at these sites is not detectable by IF (39). These findings imply that the RPA-ssDNA generated is primarily due to G-overhang elongation by following CTC1 deletion, as previously described, and not from defects in genome-wide replication or DNA damage (33). These findings are in line with the DNA fiber analysis and lack of general γH2AX staining following CTC1 removal (Figure S1C and 2C). In addition, our results confirm that RPA-ssDNA is significantly increased at telomeres in CTC1^-/-^ cells.

### Telomeric RPA-foci in CTC1^-/-^ cells are phosphorylated and γH2AX positive

Under normal conditions, excess RPA-ssDNA should lead to ATR-dependent phosphorylation of RPA at S33 (pRPA S33) (42). Based on results in Figure 2, ATR inhibition rescued G2 arrested cells following CTC1 deletion, suggesting that ATR is active, so we examined the levels of pRPA S33 in CTC1^-/-^ cells (Figure 3D-E). Similar to total chromatin-bound RPA, we observed a significant increase in pRPA S33. Previous work had also demonstrated that telomeric γH2AX was present in CTC1^-/-^ cells on metaphase spreads (33). In agreement with this study, we found that RPA-foci co-localized with γH2AX in interphase CTC1^-/-^ cells (Figure S5D). However, these foci were difficult to distinguish from background γH2AX foci in the control cells without the use of a telomere FISH probe, which precluded their detection in the global γH2AX analysis in Figure 2C. Overall, these results confirm that ATR-mediated telomeric DNA damage signaling results from CTC1 removal (33). However, while ATR is active, the signaling is not sufficient to induce global phosphorylation of CHK1 and p53.

### POT1 overexpression prevents RPA-ssDNA and rescues cell growth in CTC1^-/-^ cells

POT1 is a ssDNA binding protein and a member of the six-subunit telomere protection complex, shelterin. One of the primary roles of POT1 is to prevent ATR-dependent DNA damage signaling at telomeres through the inhibition of RPA binding to G-overhangs (43). Previous work demonstrated that POT1 overexpression (POT1-OE) partially rescues the telomeric DDR induced following CTC1 knockout (i.e. telomeric RPA and γH2AX) (33). However, POT1-OE was not able to rescue the elongated G-overhangs, indicating that POT1 is unable to substitute for CST in telomerase inhibition or C-strand fill-in. Furthermore, it suggests that endogenous POT1 levels are insufficient to outcompete RPA for the hyperextended G-overhangs caused by CTC1 deletion. To test the effects of POT1-OE on cell cycle progression and cell growth, we performed flow cytometry and growth curve analysis on CTC1^-/-^ cells overexpressing Flag-POT1 (CTC1^-/-^+POT1) (Figure 4A). In agreement with the previous study, POT-OE prevented RPA as well as pRPA S33 foci formation in the CTC1^-/-^ cells (Figure 4B and S6A). Flow cytometry and cell growth analysis revealed that not only did POT1-OE rescue G2 arrested cells but also the growth delay associated with CTC1 deletion (Figure 4C-D and S6B). These results indicate that increased telomeric RPA-ssDNA, caused by G-overhang elongation, is the primary cause of decrease proliferation following CTC1 KO and that the non-telomeric roles of CST do not significantly affect proliferation in unperturbed cells. Furthermore, they highlight the essential role of POT1 in telomere end protection and indicate that POT1 exhaustion due in an abundance of telomeric ssDNA has a profound impact on cell cycle progression and cell growth (Figure 4E).

**Figure 4:**
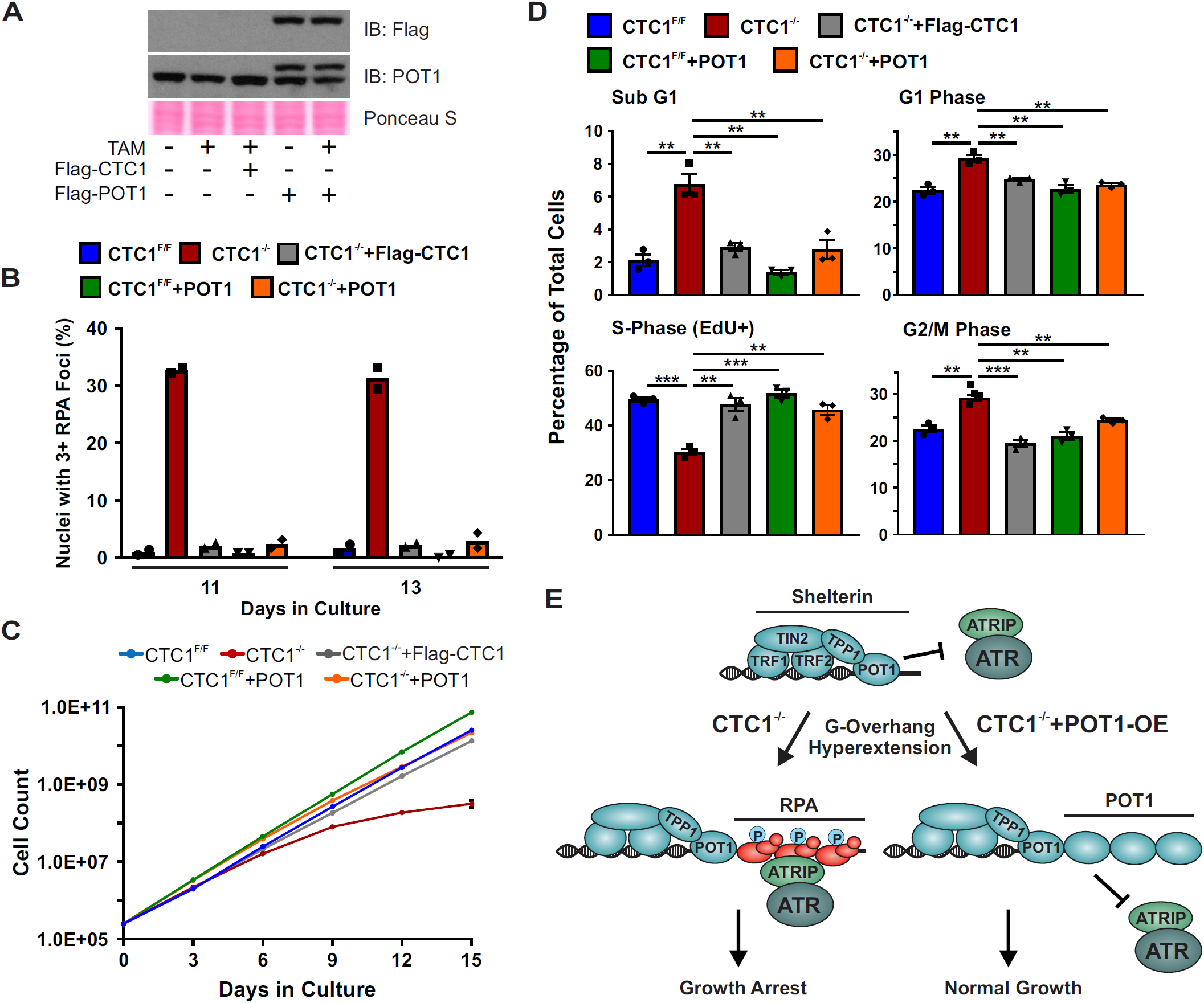
POT1 overexpression rescues cell growth and G2 arrest in the CTC1^-/-^ cells. HCT116 CTC1^F/F^+POT1 and CTC1^-/-^+POT1 cells stably express Flag-tagged POT1. **(A)** Western blot of POT1 and Flag-POT1 levels at 11 days in culture, as indicated. Ponceau S serves as a loading control. **(B)** Percentage of nuclei with three or more RPA foci 11 or 13 days after CTC1 deletion (n=2 independent biological experiments). **(C)** Representative growth curve of two independent biological replicates, as indicated. **(D)** Flow cytometry was used to assess cell cycle phases, as indicated. (n=3, independent biological replicates.) Error bars indicate the ±SEM. P-values were calculated by an unpaired, two tailed *t* test in (\**P* ≤0.05,\*\**P* ≤0.01, \*\*\**P* ≤0.001). **(E)** POT1 prevents telomeric RPA binding and thus inhibits ATR activation at telomeres. In the absence of CTC1 (CTC1^-/-^), the telomeric G-overhang is hyperextended, leading to the exhaustion of POT1 in the cell and telomeric RPA binding. This in turn leads to ATR localization and G2 arrest. However, when POT1 is overexpressed in the absence of CTC1 (CTC1^-/-^+POT1), the G-overhangs are coated by POT1, blocking RPA binding and G2 arrest.

### CTC1 KO leads to decreased levels of TopBP1 and defective ATR-mediated CHK1 signaling following exogenous replication stress

To fully activate ATR, TopBP1 is recruited by the 9-1-1 complex. Since this step is independent of ATRIP-ATR binding, we examined whether TopBP1 recruitment and protein expression were affected after CTC1 deletion. Total and chromatin-bound TopBP1 protein levels were measured at days 8, 11 and 13 following TAM addition (Figure 5A and S7A). Surprisingly, we observed a significant decrease in TopBP1 levels following conditional CTC1 KO starting at day 11 after TAM addition. These findings are significant as the timing of TopBP1 decline corresponds to the increase in RPA-ssDNA and accumulation of G2 cells following CTC1 removal. This suggested to us that ATR-mediated CHK1 signaling could be defective due to decreased cellular and chromatin-bound TopBP1. We also examined the levels of the other major ATR activator ETAA1 and found that it was also decreased in the CTC1^-/-^ cells. Since ETAA1 plays a minor role in ATR-CHK1 activation following DNA damage, it is unclear how this would affect checkpoint activation. However, it is possible that decreased ETAA1 could also contribute to changes in cell cycle progression.

**Figure 5:**
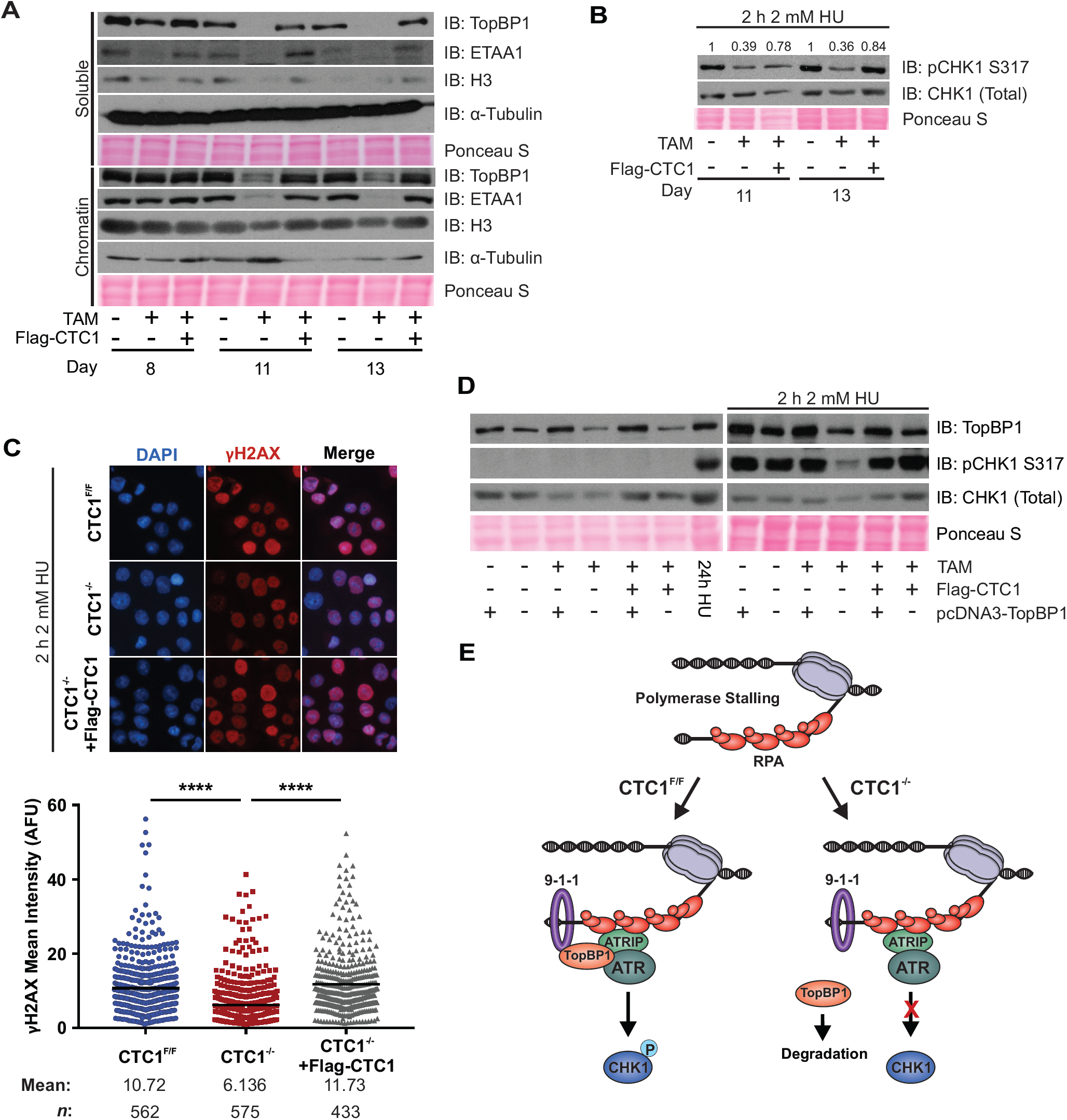
CTC1 deletion leads to decreased TopBP1 stability and inhibits ATR-CHK1 signaling following HU treatment. **(A)** Western blots of chromatin fractions from HCT116 cells at days 8, 11, and 13, as indicated. Ponceau S serves as a loading control for total protein. α-Tubulin and H3 serve as controls for the soluble and chromatin extracts, respectively. **(B)** Western blots of pCHK1 S317 and total CHK1 from whole cell lysates after treatment with 2 mM HU for 2 h in HCT116 cells, as indicated. Total CHK1 levels were used to normalize the phosphorylated CHK1 S317 signal. Quantification of signal intensity is indicated above the image. **(C)** HCT116 cells were treated with 2 mM HU for 24 hours at 13 days in culture and analyzed by immunofluorescence, as indicated. Left: Representative images of γH2AX signal. DAPI: blue, γH2AX: red. Right: Dot plots of mean γH2AX intensity per nuclei in arbitrary fluorescent units (AFU) for each cell line. Black line and numbers below the graph indicate the mean AFU (n=3, independent biological replicates). P-values were calculated by an unpaired, two tailed Mann-Whitney test (\*\*\*\**P* ≤0.0001). **(D)** Cells were transfected with pcDNA3-TopBP1 at day 8 and collected on day 11. Western blot analysis of TopBP1 and pCHK1 S317 levels in whole cell lysates. For HU treated samples, cells were treated with 2 mM HU for 2 h just prior to collection. 24h HU samples were CTC1^F/F^ cells that were treated for 24 h with 2 mM HU and were used as a control. Total CHK1 and Ponceau S serve as loading controls. **(E)** Model of CTC1 function in ATR-CHK1 activation following fork stalling. Following RPA binding, ATRIP-ATR and the 9-1-1 complex are recruited to the ssDNA damage. In the presence of CTC1 (CTC1^F/F^), TopBP1 is then recruited for ATR-CHK1 activation and G2 arrest (left). However, when CTC1 is absent (CTC1^-/-^), TopBP1 is destabilized leading to inhibition of CHK1 phosphorylation (right).

We next examined whether the decrease in TopBP1 was due to changes in gene expression. We performed qPCR to measure TopBP1 mRNA levels but did not observe any significant changes at day 8 and only minor and inconsistent changes at days 11 and 13, suggesting that decreased protein stability and not gene expression is likely responsible for the changes in TopBP1 (Figure S7B).

Since TopBP1 is important for ATR-CHK1 activation, we hypothesized that CHK1 phosphorylation might be inhibited in the presence of exogenous replication stress. To test our hypothesis, we measured CHK1 phosphorylation in cells following treatment with HU, which is known to induce global replication fork stalling, create excess ssDNA and activate the ATR-CHK1 pathway (44). Cells were treated with HU for 2 h prior to collection to generate ssDNA but not cause fork collapse and DSBs (45). Phosphorylation of CHK1 and γH2AX staining were then assessed by Western blot and IF, respectively (Figure 5B and C). When compared to control cells, we observed a significant decrease in the levels of pCHK1 S317 and γH2AX.

To see whether we could rescue pCHK1 levels, TopBP1 was exogenously expressed in the CTC1^-/-^ and control cell lines at day 8. At day 11, cells were treated with HU for 2 h and then collected for Western blot analysis. Transfection with exogenous TopBP1 resulted in similar levels of TopBP1 in the CTC1^F/F^, CTC1^-/-^ and CTC1^-/-^+Flag-CTC1 cells (Figure 5D). pCHK1 S317 was then measured as a readout of ATR activation (Figure 5D). While CHK1 phosphorylation was significantly inhibited in the CTC1^-/-^ cells, we found that the expression of TopBP1 rescued CHK1 phosphorylation in response to HU treatment. However, increasing TopBP1 did not rescue global pCHK1 S317 in CTC1^-/-^ cells in the absence of HU treatment. Overall, these results indicate that CTC1 regulates TopBP1 and is required to fully activate the ATR-CHK1 pathway in response to genome-wide replication stress (Figure 5E). These results could in part explain how CST promotes recovery from genome-wide replication stress.

## DISCUSSION

Unlike yeast, mammalian CST is involved in several distinct DNA transactions in the cell, including telomere maintenance, replication rescue, dormant origin firing, origin licensing and DSB repair (23,25-27,29,30,34). However, it has been difficult to identify the contribution of telomere versus non-telomere CST functions on genome stability and cell cycle progression. One of the major questions we sought to address in this study was the underlying cause(s) of decreased cell proliferation associated with CTC1 removal. Surprisingly, we found no significant defects in global DNA replication or increased genome-wide DNA damage. Instead, the primary cause of cell death and checkpoint activation was the accumulation of telomeric RPA due to POT1 exhaustion. Indeed, POT1 overexpression was sufficient to completely rescue cell growth and block ATR activation following CTC1 removal. Thus, telomere length regulation/protection not genome-wide replication is the primary function of CST in unstressed cells. However, this does not discount the vital non-telomeric functions of CST in rescuing stalled replication and DNA repair, which likely cause more subtle effects on genome instability following CTC1 loss. Furthermore, we determined that, while ATR is actively recruited to telomeric RPA-ssDNA, loss of CTC1 prevents global CHK1 phosphorylation. Upon further analysis, we discovered that CST regulates the ATR activator TopBP1 and that removal of CTC1 leads to a substantial decrease in CHK1 phosphorylation following global replication stress. Together, these findings highlight the complex role of human CST in maintaining genome stability. On one hand, it is necessary to prevent unwanted DNA damage signaling at telomeres. While on the other, it promotes checkpoint signaling following replication stress.

Due to their structure, telomeres must be protected from recognition as DSBs and ssDNA damage. This end protection is primarily accomplished by the shelterin complex, which blocks the DDR through inhibition of ATR, ATM and DNA-PK (46). For example, POT1 inhibits RPA binding and thus ATR activation. Deletion of POT1 in mice or chicken cells leads to G-overhang elongation, which in turn activates ATR-CHK1 signaling (47,48). This need to suppress unwanted DNA repair presents a unique situation when bona fide DNA damage occurs at telomeres. In such cases, these protection pathways must be overridden to identify and repair the damage. In the event of CTC1 deletion, telomeric RPA-ssDNA arises from elongated G-overhangs that reach kilobases in length (33). While little is known about the consequence of overextending telomere G-overhangs to such lengths, large stretches of ssDNA in the cell are quite dangerous and must be quickly repaired to prevent DNA breakage and genome instability (49). Thus, CST protects telomeres from ATR-mediated checkpoint activation and eventual cell death or senescence by inhibiting telomerase from hyperextending G-overhangs.

Despite ATR activity at RPA-bound telomeres in CTC1^-/-^ cells, our results suggest that this is insufficient to induce global CHK1 phosphorylation (Figure 2 and 3). While it is possible that low levels of pCHK1 are present but not detectable by Western blot, this seems unlikely as a significant portion of the CTC1^-/-^ cells (∼25%) contain three or more large RPA foci (Figure 3). Additionally, previous work found that replication fork stalling at a single defined repeat sequence is sufficient to induce detectable ATR-dependent CHK1 phosphorylation (Figure 3B) (50). Why CHK1 is not activated in CTC1^-/-^ cells is unclear. Yet, unlike with HU-induced fork stalling, it is not solely based on decreased levels of TopBP1 (Figure 5D). A recent study in mouse cells showed that CST can override POT1b-mediated telomere protection in S/G2, suggesting that CST may be involved in the recruitment/activation of ATR at telomeres (51). One potential mechanism to explain this inhibition, or dampening, of ATR-mediated CHK1 signaling at telomeres is that CST is required to stimulate CHK1 phosphorylation.

An interesting conundrum of ATR-CHK1 activation at elongated G-overhangs is that each telomere is predicted to contain a single loading site for the 9-1-1 complex. Yet, it is the combination of RPA/ATRIP/ATR and 9-1-1/TopBP1 that are required to activate CHK1. In order to overcome this lack of 5’-ends, CST could recruit pol α for de novo primer synthesis on RPA-bound G-overhangs, promoting additional loading of 9-1-1 complexes and TopBP1. In support of this de novo priming model, new primer synthesis was shown to stimulate checkpoint activation in *Xenopus* following fork stalling (52). In terms of shortening the extended G-overhangs, it is likely that CST/pol α are also necessary to fill-in these ssDNA regions, similar to C-strand fill-in after telomerase extension or fill-in of resected DSBs (26). Understanding how these extended G-overhangs are recognized and then “repaired” or shortened as well as understanding the interplay between POT1 and CST in ATR regulation will require additional studies. However, our work identifies CST as integral for ATR regulation at telomeres.

In regards to genome-wide ATR activation following replication stress, our results determined that regulation of TopBP1 by CST contributes to ATR-CHK1 signaling. Activation of the canonical ATR-CHK1 pathway involves five key steps: (i) binding of RPA to ssDNA, (ii) recruitment of ATRIP-ATR, (iii) loading of the 9-1-1 complex, (iv) TopBP1 interaction with RAD9 and (v) activation of ATR, leading to phosphorylation of downstream ATR targets, including CHK1 (53). Our studies provide evidence that in the absence of CTC1 steps (i-iii) remain functional but that ATR-CHK1 signaling is compromised due to decreased TopBP1 levels, short-circuiting TopBP1-dependent ATR activation (Figure 5E).

TopBP1 is integral to both checkpoint activation and DNA replication origin firing (54) so why is DNA replication and origin firing not inhibited in these cells? We propose that it is due to the decreased but not complete absence of TopBP1 (Figure 5A and S7A). In this case, while there is enough TopBP1 for origin activation, there is insufficient levels to fully activate ATR-mediated CHK1 signaling. This suggests that it may be possible to separate TopBP1 functions through regulating its protein expression. Furthermore, it implies that TopBP1 stability is regulated through multiple pathways.

While the mechanism of ATR activation by TopBP1 has been closely examined, how TopBP1 levels are regulated remains poorly understood. Previous work has shown that TopBP1 is post-transcriptionally regulated by two E3 ubiquitin ligases, UBR5 (also known as EED1 and hHYD) and HUWE1 (also known as HECTH9 and MULE) (55,56). Interestingly, TopBP1 when not chromatin associated is targeted for degradation by the ubiquitin ligase, HUWE1 (55). This protection is at least partly facilitated by the transcriptional repressor Miz1. In a similar manner, CST could stabilize TopBP1 by recruiting it to the chromatin. As to the mechanism of CST recruitment to stalled forks, we recently showed that CST interacts with the MCM2-7 helicase (25). Thus, CST could be recruited to stalled forks by MCM2-7 and then recruits TopBP1 to prevent degradation, stimulating ATR-mediated CHK1 phosphorylation. However, whether CST directly or indirectly stabilizes TopBP1 will require further studies.

In addition to TopBP1, ATR can also be activated by ETAA1. However, several studies suggest that TopBP1 is the dominant pathway for ATR-CHK1 activation following replication stress (10,57). Recent work from the Cortez lab showed that, in HCT116 cells, deletion of the AAD of ETAA1 did not inhibit ATR activation. However, attempts to delete the AAD of TopBP1 resulted in cell death, indicating that, at least in HCT116 cells, TopBP1 is the major activator of ATR (14). Thus, it is unlikely that ETAA1 could compensate for decreased TopBP1. In addition, we find that ETAA1 levels are also decreased after CTC1 KO, suggesting this potential secondary pathway is not even available for ATR-mediated CHK1 activation. Why ETAA1 levels are decreased following CTC1 deletion is unclear. However, it is possible that TopBP1 and ETAA1 are regulated through similar mechanisms.

In summary, our studies determined that, in the absence of CTC1, cells accumulate telomeric RPA-ssDNA and arrest in G2, resulting in either cell death or senescence. Intriguingly, the arrest is dependent on both ATR and RPA-ssDNA, as ATR inhibition or POT1-OE restored G2 levels (Figure 2 and 4). We also demonstrate that CTC1 promotes ATR-mediated CHK1 phosphorylation following HU-induced replication stress and that the inability to elicit an ATR-dependent checkpoint response is due to decreased levels of TopBP1. Combined, these studies establish CTC1 as a regulator of ATR-CHK1 signaling at telomeres and in response to global replication stress, which could provide a new avenue for the use of ATR/CHK1 inhibitors as cancer therapeutics.

## ACKNOWLEDGEMENT

We would like to thank Sasha Hodge, Kaury Thome and Alexander Welch for assistance with experiments and the Stewart lab members for useful discussions. We would also like to thank Dr. David Cortez for providing the ETAA1 antibody, Dr. Weei-Chen Lin for the pcDNA3-TopBP1 and Dr. Carolyn Price for the HCT116 CTC1^F/F^ cell lines and critical reading of the manuscript. This study utilized the services of the Flow Cytometry Core Facility of the COBRE Center for Targeted Therapeutics, supported by NIH grant 5P20GM109091, at the University of South Carolina with assistance from Dr. Chang-uk Lim.

