## Supplementary Materials for "Human CTC1 primarily functions in telomere maintenance/protection and promotes CHK1 phosphorylation in response to global replication stress"

### SUPPLEMENTARY MATERIALS AND METHODS

*DNA combing*

Cells were labeled with IdU (50 μM) for 30 min, washed three times with 1x PBS and then labeled with CldU (100 μM) for 30 min. Cells were then collected, washed once with 1x PBS and diluted to ~3,300/μl. Agarose plugs were then made and prepared for DNA combing, as previously described (63). DNA fibers were combed on silanized coverslips according to the manufacturer's instructions (Genomic Vision). Coverslips were then baked at 80°C for 2 h, washed once with 1x PBS and denatured with 0.5 M NaOH/1 M NaCl for 8 min. Following two 1x PBS washes, the coverslips were blocked in 3% BSA/1x PBS for 30 min followed by incubation with two α-BrdU antibodies (Accurate Chemical [OBT0030] and BD [347580]) (1:100), each with specificity to either IdU or CldU, for 2 h at 37°C. After three PBST washes, goat α-mouse AlexaFluor 594 and goat α-rat AlexaFluor 488 (1:500) secondary antibodies were incubated on the coverslips for 1 h at 37°C. Coverslips were washed three times with 1x PBST, dehydrated and mounted on slides with mounting media. Images were then taken under 40x on an EVOS epifluorescence microscope (Thermo) and scored using ImageJ.

*Caspase-Glo 3/7 Assay*

Active apoptosis was measured using the Promega Caspase-Glo 3/7 Assay (G8090), according to the manufacturer's instructions. DMSO or 5 μM VE-821 was added for 24 h, as indicated. Each treatment was produced in triplicate and at least two independent, biological replicates performed. The luciferase activity was measured on a Spectra Max i3 (Molecular Devices) and analyzed by Softmax Pro 6 (Molecular Devices).

*RPA Flow Cytometry*

The cells were collected and washed with 1x PBS. After the supernatant was removed, the cell pellets were resuspended in 500 μl 1x CSK buffer (10 mM HEPES pH 7.4, 300 mM sucrose, 100 mM NaCl, and 3 mM $MgCl_2$ diluted in dd$H_2O$) with 0.1% Triton-X and incubated at room temperature for 5 minutes. Then, with gentle vortexing, 5 mL of ice-cold 100% methanol was added to the tubes drop wise. Tubes were then placed at -20ºC for 10 min. Then 5 mL of 1% BSA-PBS was added to the samples followed by centrifugation at 2000 *x g* for 5 minutes. The supernatant was removed and the cell pellets were resuspended in 5 mL 1% BSA-PBS. Samples were then stored at 4ºC at least overnight.

For analysis of chromatin-bound RPA, cells were then centrifuged at 2000 *x g* for 5 minutes and the supernatant removed. The cell pellets were then resuspended in 5 mL of 1% BSA-PBST and incubated at room temperature for 15 minutes. Samples were spun down at 1000 x *g* for 5 minutes. The supernatant was removed and the cell pellet was resuspended in 200 µL of RPA32 primary antibody diluted in 1% BSA-PBST (1:500). The samples were incubated with the primary antibody for at least 1 h at RT with mild vortexing halfway through incubation. 5 mL of 1% BSA-PBST was then added. Samples were spun down at 1000 x *g* for 5 minutes and the supernatant was removed. Cells were then resuspended and incubated in 200 µL of goat α-mouse AlexaFluor 647 (1:500) in 1% BSA-PBST for at least 1 h at RT, protected from light, with mild vortexing halfway through incubation. Afterwards, 5 mL of 1% BSA-PBST was added and samples were spun down at 1000 x *g* for 5 minutes and the supernatant was removed.

### SUPPLEMENTARY FIGURES

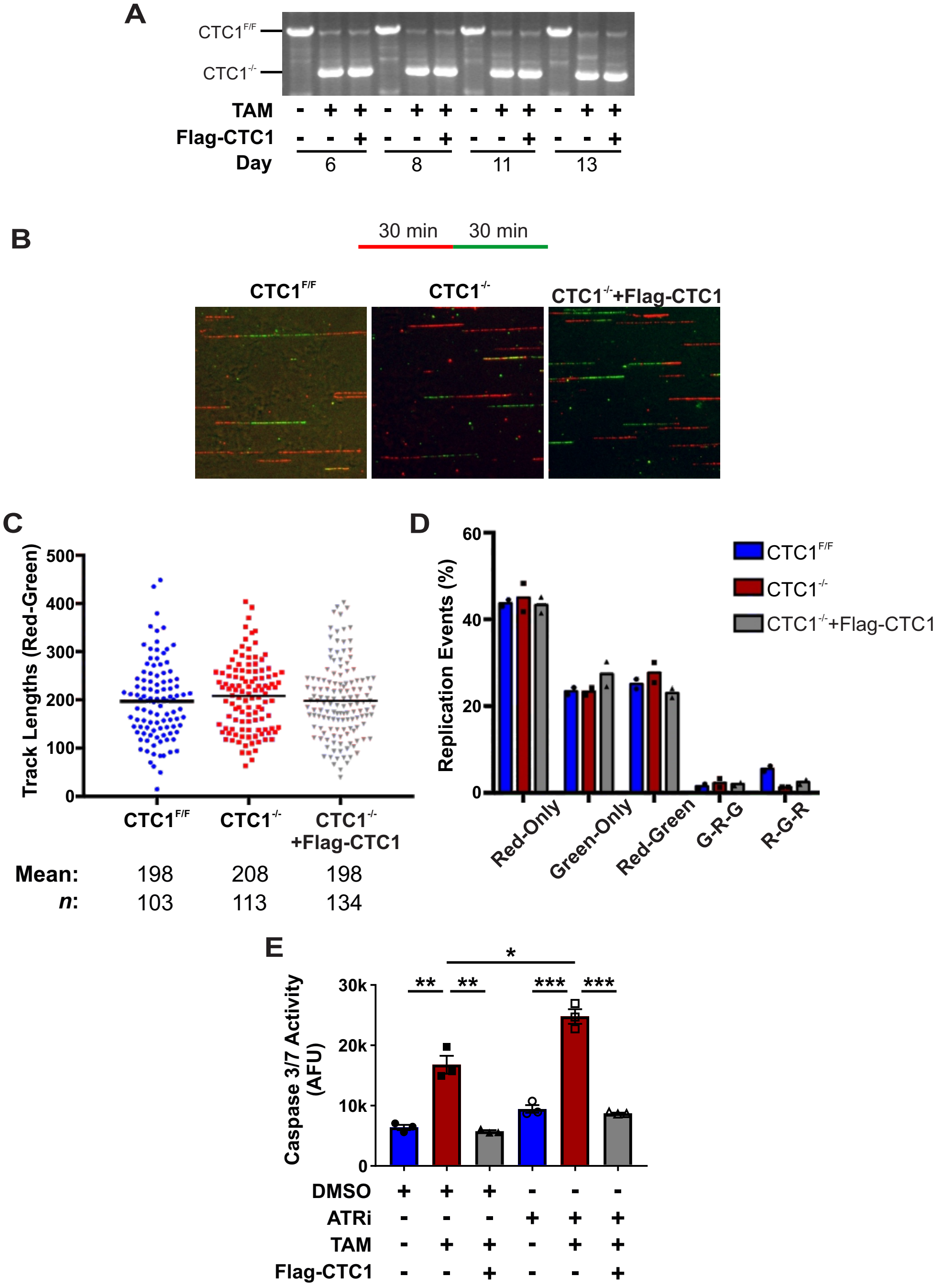


**Figure S1: Analysis of CTC1 gene disruption, DNA combing and apoptosis in CTC1$^{-/-}$ cells. (A)** Agarose gel of truncated CTC1 PCR product in CTC1$^{-/-}$ and CTC1$^{-/-}$+Flag-CTC1 samples. **(B-D)** DNA combing analysis in the HCT116 cells, as indicated. Cells were collected and processed 11 days after TAM addition. (n=2, independent biological experiments) **(B)** Images are representative of fibers staining. **(C)** Dot plot of track length for elongating forks (Red-Green). Black line and numbers below the graph indicate the mean length in arbitrary fluorescence units (AFU). *n* indicates the number of total tracks scored. **(D)** Percentage of replication events. Red-only: stalls or terminations, Green-only: origins fired during second label (CldU), Red-Green: elongating forks, G-R-G (Green-Red-Green): origins fired in first label (IdU), R-G-R (Red-Green-Red): terminations. Number of events scored: CTC1$^{F/F}$: 451, CTC1$^{-/-}$: 393, CTC1$^{-/-}$+Flag-CTC1: 572. **(E)** Fold change of apoptosis as measured by the Caspase-Glo 3/7 assay. The ATR inhibitor (5 µM, VE-821) was added for 24 h prior to measurement. (n=3, independent biological experiments). Error bars indicate the ±SEM. P-values were calculated by an unpaired, two tailed *t* test in (**P* ≤0.05, ***P* ≤0.01, ****P* ≤0.001).

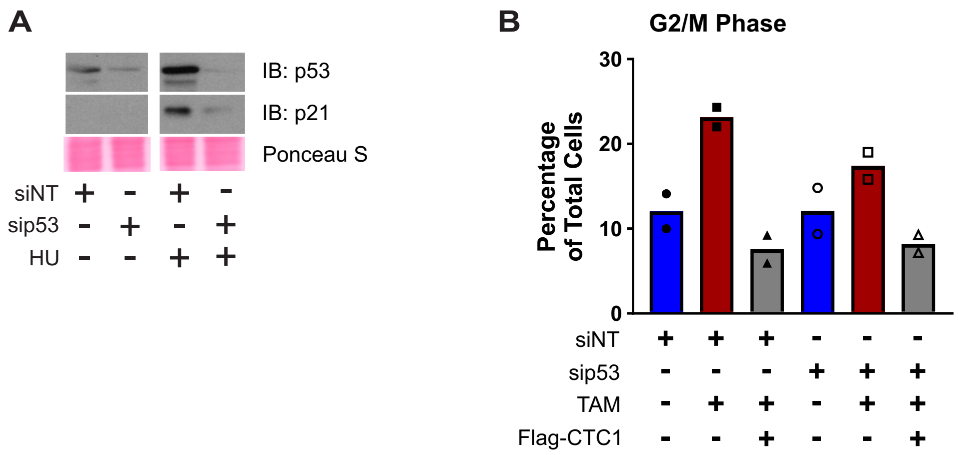


**Figure S2: p53 knockdown rescues G2 arrested cells in CTC1$^{-/-}$ cells.** Cells were treated with either 5 nM of siRNA to a non-targeting sequence (siNT) or 5 nM siRNA to p53 (sip53) for 72 hours. **(A)** Western blot analysis of 5 nM sip53 knockdown efficiency in untreated and 2 h 2 mM HU treated samples. Ponceau S is used as a loading control. **(B)** Cells were labeled with EdU, fixed, and flow cytometry performed to analyze cell cycle profile following treatment with sip53. Graph represents the number of G2/M cells from two independent biological experiments, as indicated.

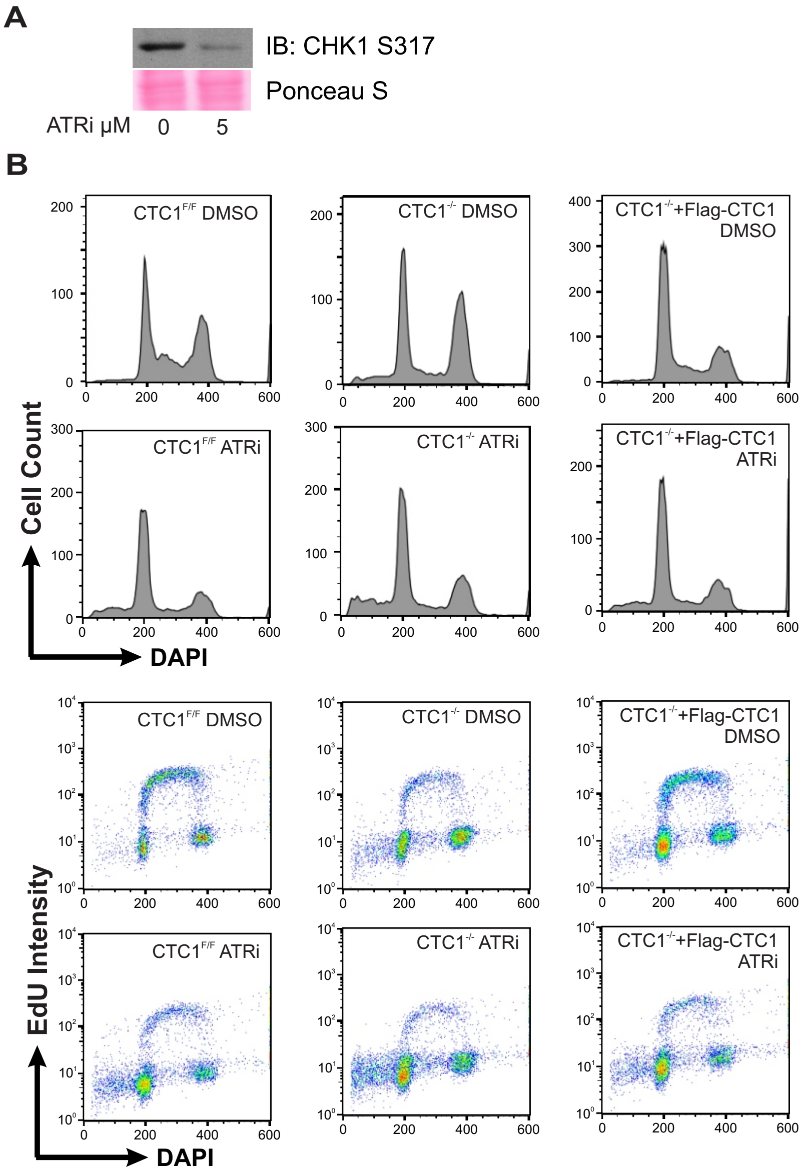


**Figure S3: ATR inhibition rescues G2 arrested cells following CTC1 deletion. (A)** Western blot analysis of HCT116 CTC1$^{F/F}$ cells treated with 5 µM VE-821 (ATR inhibitor) for 24 h. 2 mM HU was added 2 h prior to collection to induce CHK1 S317 phosphorylation. Ponceau S is used as a loading control. **(B)** Cells were treated with either DMSO or 5 µM VE-821 (ATRi) for 24 hours. Top: DAPI versus cell count. Bottom: DAPI versus EdU signal intensity.

CTC1$^{-/-}$ Day 13 (DAPI/Telomere/RPA)

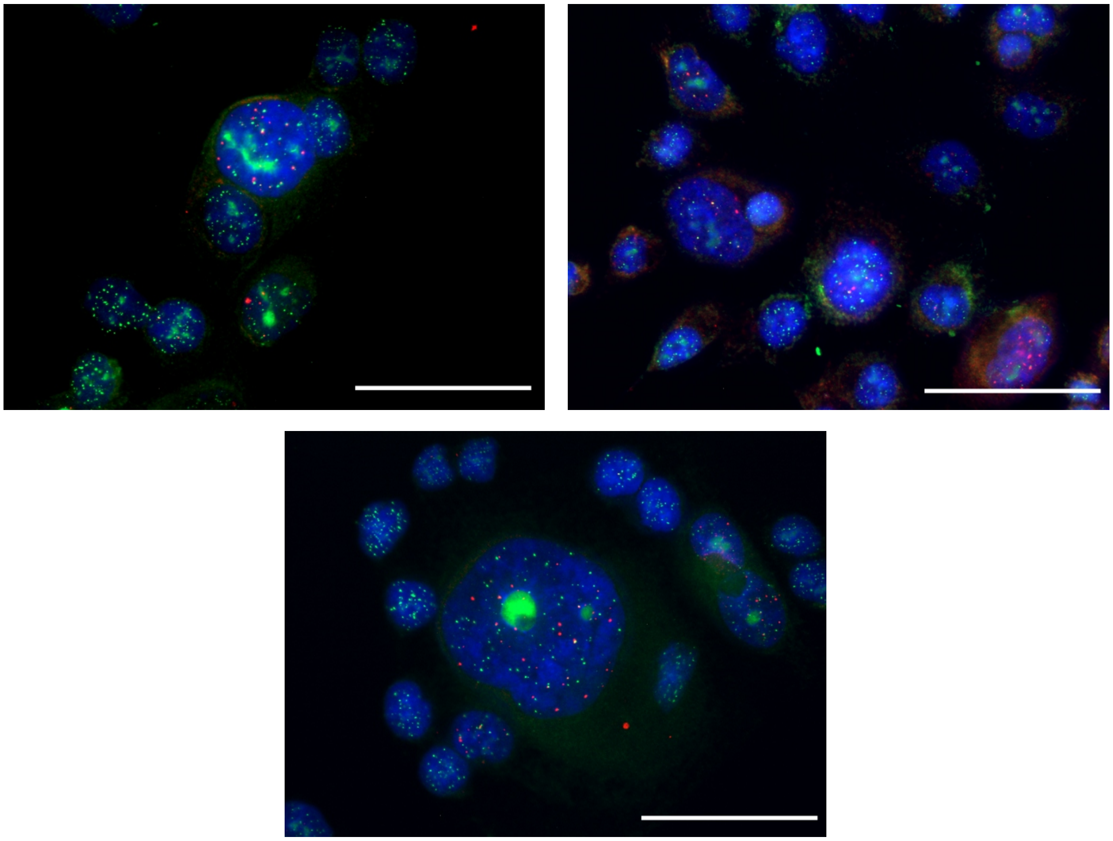

**Figure S4:** Representative images of CTC1$^{-/-}$ cells at day 13 after TAM addition. Scale bar represents 50 µm.

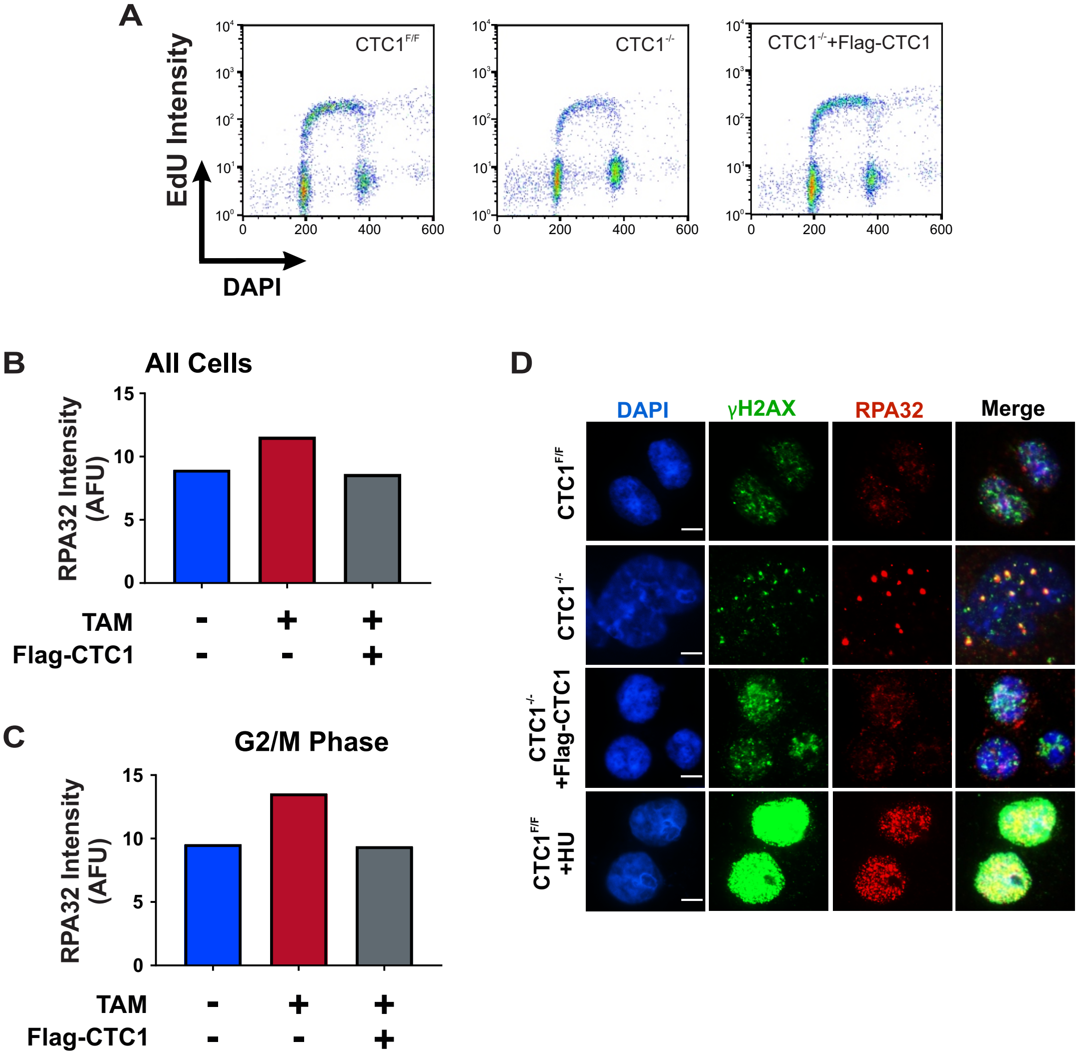


**Figure S5: RPA levels are increased in CTC1$^{-/-}$ cells and co-localize with γH2AX. (A-B)** Flow cytometry analysis of chromatin-bound RPA32 in pre-extracted HCT116 cells, as indicated. **(A)** DAPI versus EdU signal. **(B)** Graph of RPA32 signal across all cells. **(C)** RPA32 intensity in G2/M cells. **(D)** Representative images of co-localization of RPA and γH2AX foci in HCT116 cells, as indicated. CTC1$^{-/-}$ +HU samples were treated with HU for 24 h prior to fixation and is used as a control to show the intensity of γH2AX foci in the presence of global replication stress.

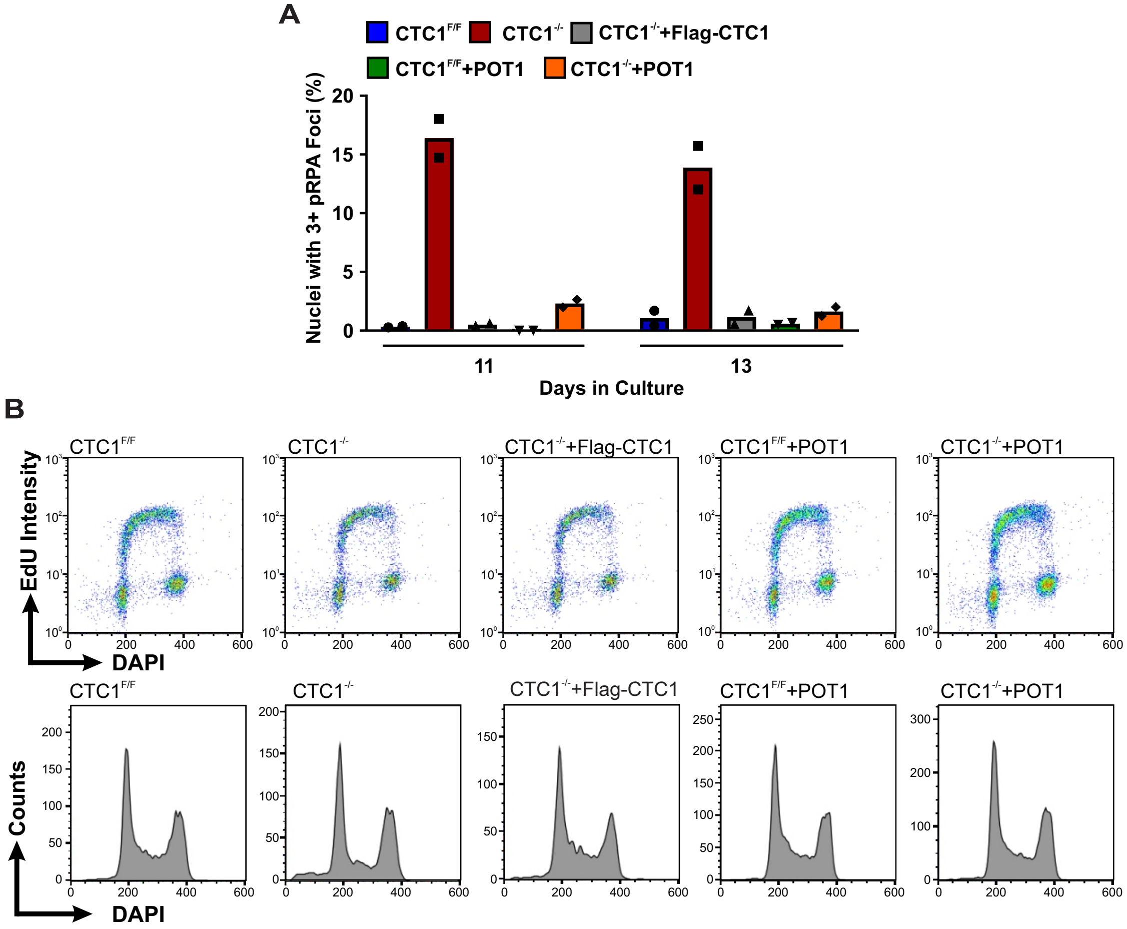


**Figure S6: Overexpression of POT1 prevents pRPA S33 and G2 arrest in CTC1$^{-/-}$ cells. (A)** Percentage of nuclei with three or more pRPA S33 foci 11 or 13 days after CTC1 deletion (n=2 independent biological experiments). **(B)** HCT116 cells were labeled with EdU for 30 min, collected and run by flow cytometry, as indicated. Top: DAPI versus EdU signal intensity. Bottom: DAPI versus cell count.

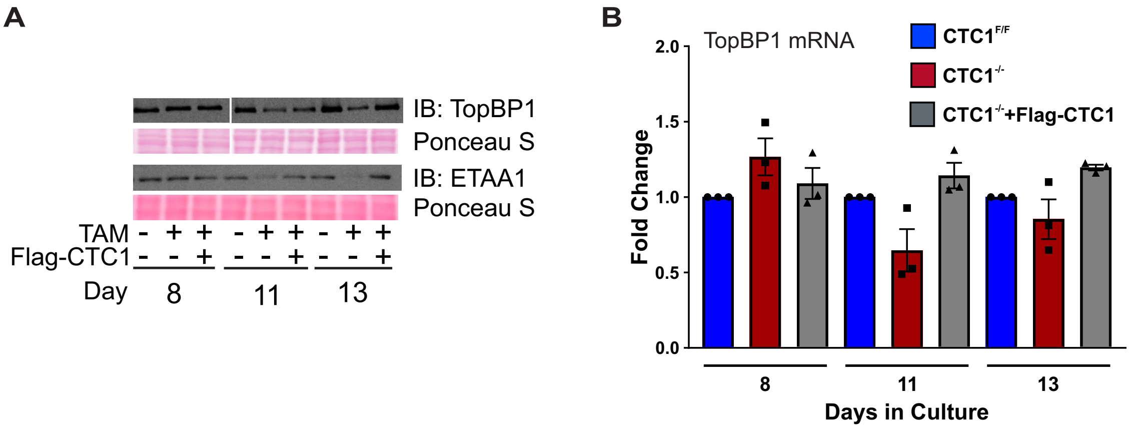


**Figure S7: Total TopBP1 and ETAA1 levels are decreased and TopBP1 mRNA levels are not significantly altered after CTC1 removal. (A)** Western blot analysis of whole cell lysates from HCT116 cells, as indicated. Ponceau S is used as a loading control. **(B)** Graph of the fold change in TopBP1 mRNA levels at day 8, 11, and 13 following TAM addition in HCT116 cells. CTC1$^{-/-}$, and CTC1$^{-/-}$ +Flag-CTC1 fold change is normalized to the CTC1$^{F/F}$ sample on each day. (n=3 three independent biological experiments.)
